# Loop extrusion as a mechanism for DNA Double-Strand Breaks repair foci formation

**DOI:** 10.1101/2020.02.12.945311

**Authors:** Coline Arnould, Vincent Rocher, Thomas Clouaire, Pierre Caron, Philippe. E. Mangeot, Emiliano. P. Ricci, Raphael Mourad, Daan Noordermeer, Gaëlle Legube

**Affiliations:** LBCMCP, Centre de Biologie Intégrative (CBI), CNRS, Université de Toulouse, UT3; CIRI – International Center for Infectiology Research, Inserm, U1111, Université Claude Bernard Lyon 1, CNRS, UMR5308, Ecole Normale Supérieure de Lyon, Univ Lyon, F-69007, Lyon, France; Laboratoire de Biologie et Modélisation de la Cellule, Université de Lyon, INSERM U1210, CNRS UMR 5239, Ecole Normale Supérieure de Lyon, Université Claude Bernard Lyon 1, F-69007 Lyon, France; Université Paris-Saclay, CEA, CNRS, Institute for Integrative Biology of the Cell (I2BC), Gif-sur-Yvette, France

**Keywords:** Cohesin, Loop extrusion, DNA Double-Strand Breaks, γH2AX, DNA Damage Response, Topologically Associated Domains

## Abstract

DNA Double-Strand Breaks (DSBs) repair is essential to safeguard genome integrity. Upon DSBs, the ATM PI3K kinase rapidly triggers the establishment of megabase-sized, γH2AX-decorated chromatin domains which further act as seeds for the formation of DNA Damage Response (DDR) foci^1^. How these foci are rapidly assembled in order to establish a “repair-prone” environment within the nucleus is yet unclear. Topologically Associating Domains (TADs) are a key feature of 3D genome organization that regulate transcription and replication, but little is known about their contribution to DNA repair processes^2,3^. Here we found that TADs are functional units of the DDR, instrumental for the correct establishment of γH2AX/53BP1 chromatin domains in a manner that involves one-sided cohesin-mediated loop extrusion on both sides of the DSB. We propose a model whereby H2AX-containing nucleosomes are rapidly phosphorylated as they actively pass by DSB-anchored cohesin. Our work highlights the critical impact of chromosome conformation in the maintenance of genome integrity and provides the first example of a chromatin modification established by loop extrusion.

---

DNA double-strand breaks induce the formation of DDR foci, which are microscopically visible and characterized by specific chromatin modifications (γH2AX, ubiquitin accumulation, histone H1 depletion) and the accumulation of DDR factors (53BP1, MDC1)^4–6^. Previous evidence indicated that chromosome architecture may control γH2AX spreading. Indeed, γH2AX domain boundaries were found in some instances to coincide with TAD boundaries^7^. Moreover, super-resolution light microscopy revealed that CTCF, which demarcates TAD boundaries in undamaged cells, is juxtaposed to γH2AX foci^8^. In addition, 53BP1 can form nanodomains which frequently overlap with a TAD, as detected by DNA-FISH^9^. High-resolution ChIP-seq mapping following the induction of multiple DSB at annotated positions (using human DIvA cells)^10^ revealed that the spreading of these DDR foci components on nearby chromatin follows a highly stereotyped pattern (one example shown in Fig 1a)^5^. We hypothesized that such pattern could be governed by pre-existing high-order chromatin structure established prior to DSB induction. In order to relate the spreading of DDR foci components with chromosome conformation, we performed high-resolution 4C-seq experiments in undamaged human DIvA cells. As viewpoints we selected three genomic locations that are damaged in DIvA cells following activation of the AsiSI restriction enzyme as well as one undamaged control region. The chromatin conformation around these three viewpoints in normal (undamaged) condition was remarkably similar to the distribution of γH2AX determined post DSB induction (Fig. 1a-b, Fig. S1a), suggesting that initial chromosome architecture dictates γH2AX spreading and downstream events such as accumulation of MDC1, ubiquitin and 53BP1 following DSB. To prove that DDR domains do not spread into neighboring self-interacting domains, we focused on an AsiSI-induced DSB located on chr1, for which spreading of DDR foci components is profoundly asymmetrical (Fig. 1b, red signal). 4C-seq performed at two independent viewpoints separated by 470 kb revealed the existence of two adjacent self-interacting domains with a boundary corresponding to the abrupt drop in γH2AX (Fig. 1c, blue signal, TAD boundary is indicated by the dotted line). This strongly suggests that pre-existing chromatin domains, established before any damage occurs, constrain the spread of DDR foci. To generalize this finding, we performed Hi-C in undamaged DIvA cells. Strikingly, computed Topologically Associating Domains (TADs) boundaries coincided with a sharp decrease in γH2AX signals (Fig. 1d-e). In agreement, γH2AX, MDC1 and 53BP1 were significantly more enriched within the damaged TADs compared to neighboring TADs (Fig. S1b), although spreading through boundaries was observed to some extent, in agreement with the rather moderate insulation properties of these domains and boundaries^11^.

**Figure 1.**
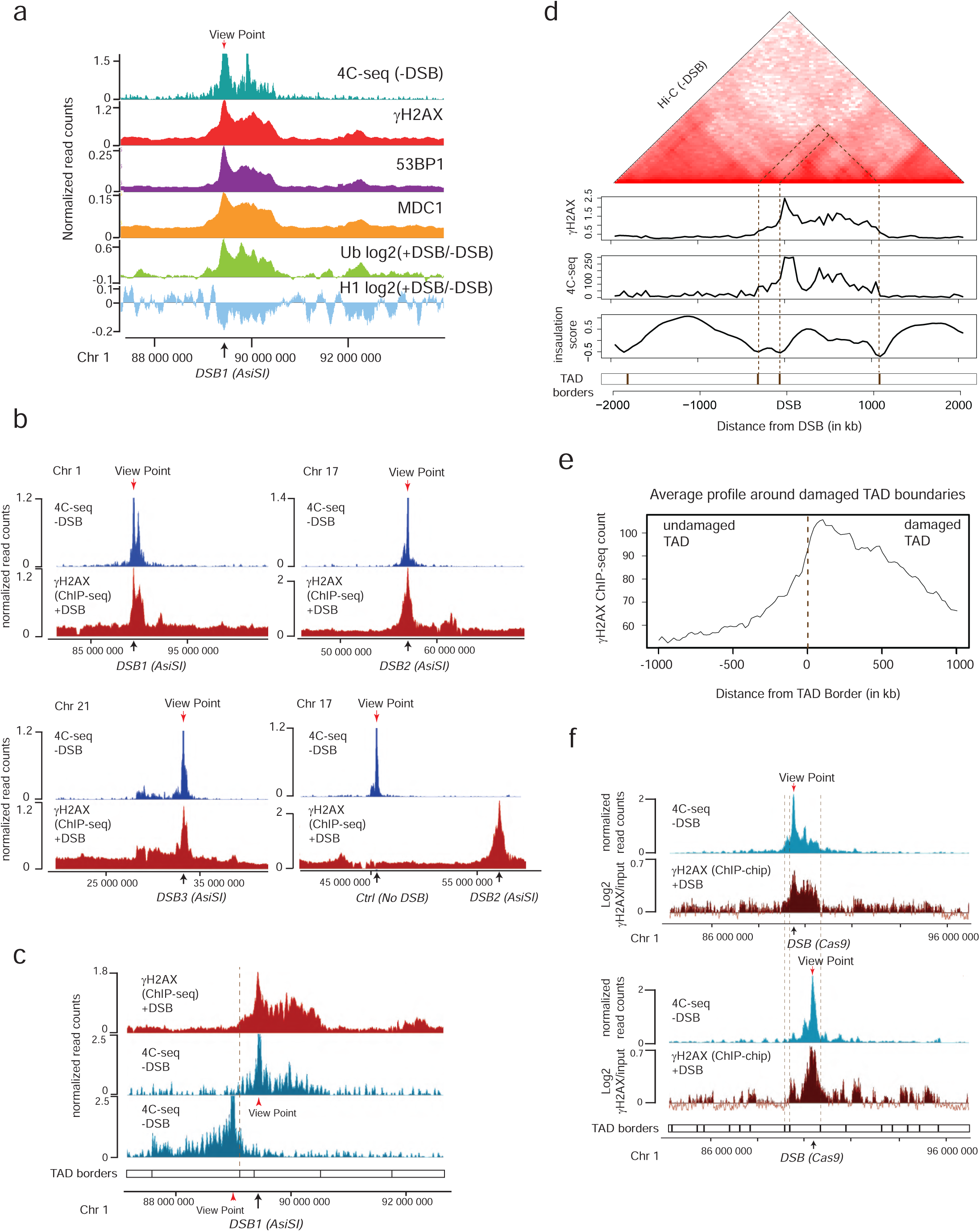
TADs are functional units governing DDR chromatin domains establishment. **(a)** Genome browser screenshot representing the 4C-seq signal in undamaged cells (-DSB), the histone H1 and Ubiquitin (FK2) ChIP-seq log2 ratio in damaged versus undamaged condition (+DSB/-DSB) and the γH2AX, MDC1 and 53BP1 ChIP-seq signals after DSB induction (+DSB) as indicated. ChIP-seq data are smoothed using 50kb span, 4C-seq using a 10kb span. The AsiSI site is indicated by an arrow. **(b)** Genome browser screenshots representing the 4C-seq signal before DSB induction (-DSB) and the γH2AX ChIP-seq signal after DSB induction (+DSB) for different viewpoints localized at three AsiSI sites or a Control region. One representative experiment is shown. ChIP-seq and 4C-seq data are smoothed using a 50kb span. The AsiSI sites are indicated by arrows. **(c)** Genome browser screenshot representing the γH2AX ChIP-seq signal after DSB induction (+DSB) and the 4C-seq signal before DSB induction (-DSB) for two viewpoints localized at the AsiSI site or 470 kb upstream of the AsiSI site. Viewpoints are indicated by red arrows, the DSB by a black arrow. ChIP-seq and 4C-seq data are smoothed using 10 kb span. **(d)** Hi-C contact matrix of a region of the chromosome 1 in DIvA cells before DSB induction (top panel). γH2AX ChIP-seq after DSB induction, 4C-seq signal, insulation score and TAD borders before DSB induction are shown. **(e)** Average profile of γH2AX ChIP-seq after DSB induction centered on the closest TAD border to the 174 best-induced DSBs (damaged TAD on the right). **(f)** Genome browser screenshots representing the 4C-seq signal (smoothed using a 10 kb span) before DSB induction (-DSB) (in blue) using view points as indicated (red arrows). γH2AX ChIP-chip (log2 sample/input, smoothed using 500 probes span) after DSB induction with CRISPR/Cas9 (black arrows) are shown in red.

To further investigate if TADs dictate γH2AX spreading, we used the CRISPR/Cas9 system to induce a single DSB at two designed positions within the same TAD, and investigated both chromosome conformation and γH2AX distribution. Cas9-induced DSB recapitulated the γH2AX spreading observed when inducing a DSB at the same genomic location by AsiSI (Fig. S1c), thus confirming that γH2AX spreading is independent of the DSB induction method. Moving the DSB to a further downstream position in the TAD triggered a change in the γH2AX profile that was notably similar to the 3D interaction pattern of this genomic region, yet it remained constrained within the same TAD (Fig. 1f). Altogether these data indicate that pre-existing self-interacting domains facilitate and demarcate the formation of γH2AX domains. Given that γH2AX is seeding further signaling events leading to the stable assembly of DDR foci and full checkpoint activation, this indicates that genome organization within TADs is critical for the response to DNA damage.

In order to gain more insights into the mechanism that mediates γH2AX establishment on entire self-interacting domains, we further profiled ATM which is the main DDR kinase that catalyzes H2AX phosphorylation upon damage in DIvA cells^12^. Activated ATM (autophosphorylated on S1981) binding was restricted to the immediate vicinity of the DSB (< 5kb) in sharp contrast with the pattern observed for γH2AX (see Fig. 2a for one example and Fig. 2b for the average profiles around the eighty best-induced DSBs). This indicates that H2AX phosphorylation is not mediated by the linear spreading of the kinase on entire TADs.

**Figure 2.**
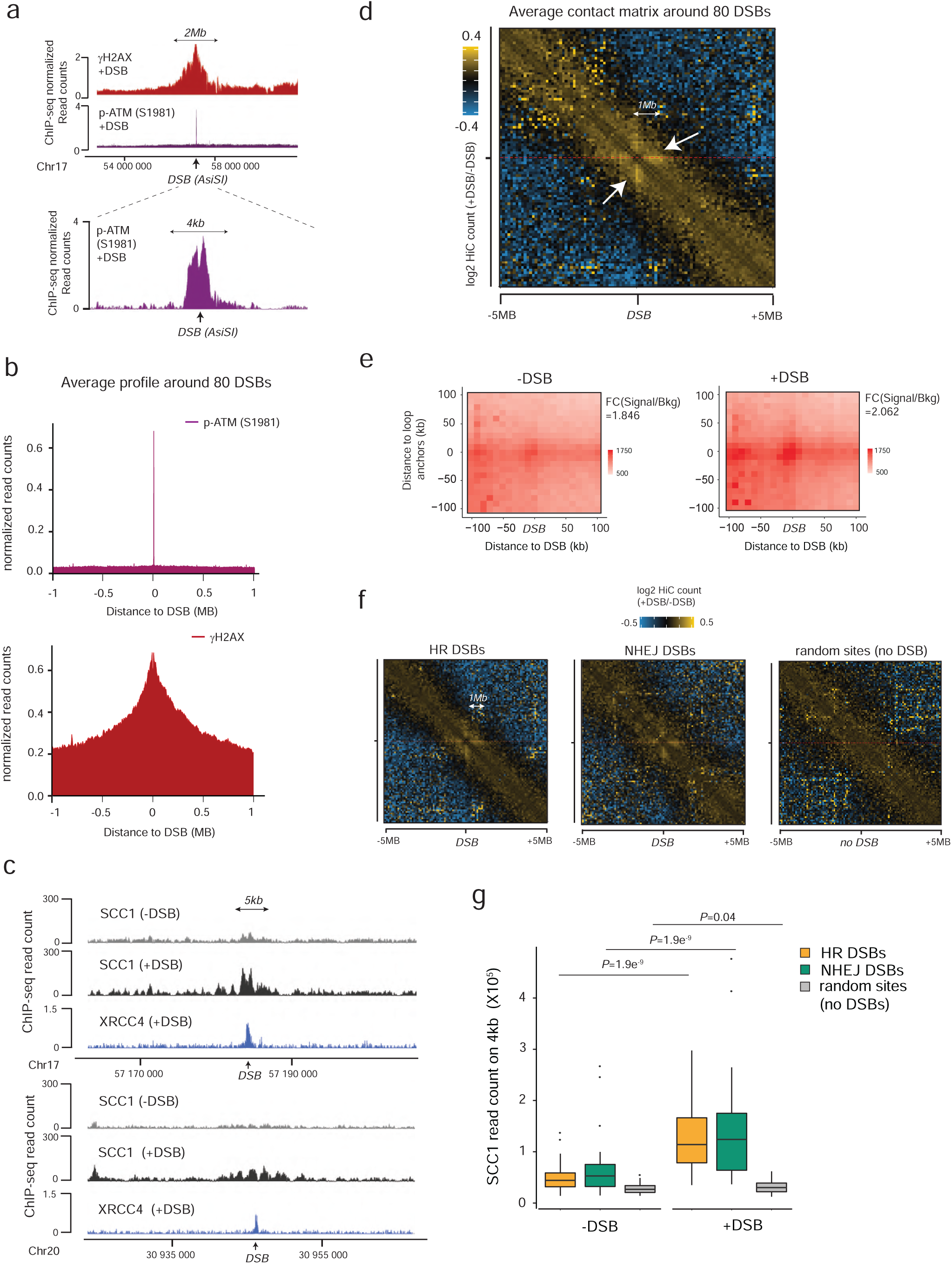
DSB-anchored cohesin mediates loop extrusion. **(a)** Genome browser screenshot representing γH2AX and p-ATM (S1981) ChIP-seq signals after DSB induction (+DSB) on an 8 Mb window (top panel) and a 15 kb window (bottom panel) around an AsiSI site (black arrow). **(b)** Average profile of p-ATM (S1981) (top panel) and γH2AX (bottom panel) ChIP-seq on a 2 Mb window around the eighty best-cleaved DSBs in DIvA cells. **(c)** Genome browser screenshots representing the calibrated SCC1 ChIP-seq before (-DSB) and after DSB induction (+DSB). XRCC4 (an NHEJ component) ChIP-seq after DSB induction (+DSB) is also shown. DSB are indicated by black arrows. **(d)** Averaged Hi-C contact matrix representing the log2 ratio between damaged versus undamaged cells (from 2 biological replicates) centered on the eighty best-induced DSBs (100 kb resolution on a 10 Mb window). Stripes are indicated by white arrows. **(e)** Aggregate Peak Analysis (APA) plot on a 200kb window (10 kb resolution) before (-DSB) and after DSB induction (+DSB), calculated between the DSBs and loops anchors (n=525 pairs). The fold change between the signal (central pixel) and the background (upper left corner 5×5 pixels) is indicated. **(f)** Averaged Hi-C contact matrix representing the log2 ratio between damaged versus undamaged cells (from 2 biological replicates) around 30 DSBs found as repaired by HR (left panel), or by NHEJ (mid panel) 5 as well as around 30 random undamaged sites (right panel), using a100 kb resolution on a 10 Mb window. **(g)** Boxplots representing the SCC1 ChIP-seq enrichment before and after DSB induction as indicated, on 4 kb around DSBs repaired by HR (yellow) or by NHEJ (green). Values for random undamaged sites (grey) are also shown. P values are indicated (paired two-sided Wilcoxon test).

Proper TAD structure requires the active involvement of the ring-shaped cohesin protein complex^13,14^, which was initially identified for its essential role in sister chromatid cohesion. Of importance, strong evidence support a role of cohesin in the maintenance of genome integrity^15,16^ and cohesin accumulates at sites of damage, which may be in line with a role in sister chromatid cohesion during Homologous Recombination (HR) in S/G2^17–23^. Yet, cohesin enrichment has been identified at DSBs throughout the cell cycle as well, which argues against an exclusive role for cohesin in HR^7,16^. In order to get insights in the cohesin binding status at DSBs at high resolution, we performed calibrated ChIP-seq profiling of the SCC1 cohesin subunit, in both undamaged and damaged conditions. Notably, cohesin was enriched at sites of damage spanning 2-5kb around the DSB (Fig. 2c).

Cohesin has been proposed to structure TADs by an active, ATP-dependent, loop extrusion mechanism^24–27^. In this model, once loaded onto chromatin, cohesin leads to the formation and processive enlargement of DNA loops, eventually arresting at boundary elements. Increased cohesin around DSBs could thus indicate ongoing loop extrusion at site of damage. We hence set to analyze 3D genome organization by Hi-C before and after DSB induction in DIvA cells, focusing on *cis* interaction frequencies around DSBs. Interestingly, within the Hi-C map, a unique pattern of “stripes” emerges from both sides of the DSBs (Fig. 2d, white arrows). Single stripes were previously reported as the characteristic of one-sided loop extrusion, where localized cohesin loading or fixation allows a locus to act as a loop anchor with increased interaction with an entire neighboring domain^27–29^. Therefore, we performed Aggregate Plot Analysis (APA) to assess looping between the DSB position and neighboring anchors. Notably, the APA score increased following production of DSBs (Fig. 2e) indicating that the DSBs themselves displays loop anchoring properties.

We previously determined which repair pathway (*i*.*e*. Homologous Recombination (HR) or Non Homologous End Joining (NHEJ)) is preferentially utilized at each DSB induced by AsiSI in DIvA cells^30^. Importantly, an equivalent stripe pattern was observed at both DSBs repaired by HR or NHEJ (Fig. 2f). In agreement, SCC1 accumulates on a 4 kb window around DSBs irrespective of the pathway used for repair (Fig. 2g).

Altogether these data suggest that cohesin accumulates at either side of a DSB, irrespective of the pathway used for repair, to induce a divergent one-sided loop extrusion towards (and thus increased contacts with) the surrounding regions on both sides of the break.

In order to further investigate DSB-anchored loop extrusion, we performed 4C-seq before and after DSBs induction, using viewpoints located at the exact positions of three DSBs induced in DIvA cells (same viewpoints as in Figure 1). Importantly, the overall TADs structure and boundaries were well maintained post DSB induction (Fig. 3a, Fig. S2a) indicating that chromosome conformation within TAD is not completely reshuffled upon damage induction on the genome. Yet, as expected from Hi-C data, we could detect increased interactions between viewpoints and surrounding loci post-DSB (Fig. 3b, 3c, Fig. S2b), which was not the case when using a control undamaged sequence as a viewpoint (Fig. 3c, Fig. S2b). If DSB-anchored, cohesin-mediated loop extrusion is responsible for enhanced interaction frequency of the DSB with neighboring sequences post DSB induction, such behavior should be abolished following cohesin depletion. Indeed, depletion of SCC1 (Fig. S2c-d) strongly impaired the overall increase in contacts between the DSBs and its neighboring sequences in damaged TADs (Fig. 3d-e). This data indicates that the ability of the DSB to contact neighboring loci within the damaged TAD is a proper DNA Damage Response and cannot solely be explained by the physical disruption of the DNA. Importantly, it depends on the cohesin complex, in agreement with a DSB-anchored loop extrusion mechanism.

**Figure 3.**
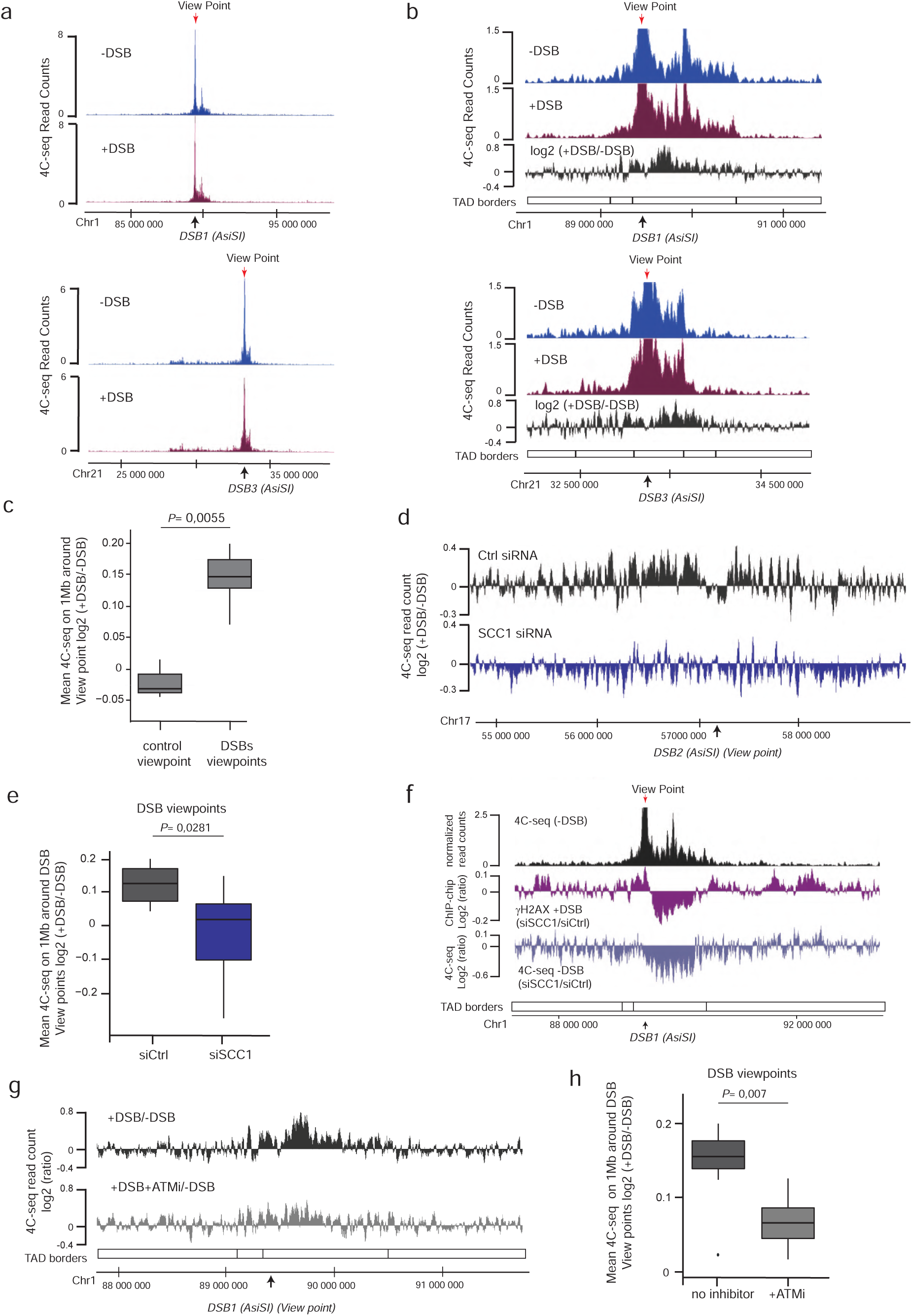
Increased interactions between DSBs and cis loci depends on ATM and cohesin. **(a)** Genome browser screenshots representing the 4C-seq signals on a ∼15 Mb window (smoothed using a 10 kb span) before (blue) and after (purple) DSB induction as indicated using two independent DSB viewpoints. One representative experiment is shown. **(b)** Same as in (a) but zoomed on a ∼3 Mb window. Differential 4C-seq (Log2 +DSB/-DSB) is also indicated (black). **(c)** Differential 4C-seq signal (Log2 +DSB/-DSB) was computed on 1 Mb around four independent viewpoints located at DSBs (“DSBs viewpoints”) and one control region (“Control viewpoint”) and across four independent biological replicates. P value between control and DSBs viewpoints is indicated (two-sided Wilcoxon test). **(d)** Genome browser screenshot representing differential 4C-seq signal (Log 2 +DSB/-DSB, 10 kb smoothed) in control siRNA condition (black) or in SCC1 siRNA condition (blue). **(e)** Boxplot representing cis interactions computed as in (c) for four DSBs viewpoints (two independent biological experiments) upon control or SCC1 depletion by siRNA as indicated. P value between control and SCC1 siRNA is indicated (two-sided Wilcoxon test). **(f)** Genome browser screenshot representing 4C-seq before DSB induction using a DSB viewpoint (DSB is indicated by a black arrow, Viewpoint by a red arrow). Purple (middle) track represents the differential γH2AX signal obtained after DSB induction by ChIP-chip in SCC1-depleted versus control cells (expressed as the γH2AX log2 ratio siSCC1/siCtrl). Light blue track (bottom track) shows the differential 4C-seq signal obtained in SCC1-depleted versus control cells before DSB induction (log2 siSCC1/siCtrl). **(g)** Genome browser screenshot representing differential 4C-seq signal (Log 2 +DSB/-DSB, 10kb smoothed) in control condition (black) or upon ATM inhibition (grey). **(h)** Boxplot representing cis interactions computed as in (c) for four DSBs viewpoints (three biological experiments) in control condition or upon ATM inhibition as indicated. P value between control and ATM inhibited cells is indicated (two-sided Wilcoxon test).

In order to investigate whether cohesin depletion, which triggers decreased interaction between the DSB and its neighboring sequences, also results in impaired DDR foci establishment, we further compared γH2AX spreading and the 4C-seq profile obtained in cells depleted for the SCC1 cohesin subunit^7^. Notably, γH2AX spreading was impaired at genomic loci showing decreased *cis* contacts, compared to control, SCC1-proficient, cells (Fig. 3f).

Of interest, NIPBL, recently found to be necessary for loop extrusion^24,25^, is recruited at laser induced damages in a manner that depends on DDR kinases^31^. We therefore assessed the consequences of pharmaceutical inhibition of the ATM kinase activity on the interaction frequency post-DSB. As expected from previous work^12^, ATM inhibition nearly abolished γH2AX foci formation after DSBs (Fig. S2e). Notably, upon ATM inhibition the ability of the DSB to engage in contacts with proximal sequences within damaged TADs was almost completely lost (Fig. 3g-h).

Taken together these data suggest that cohesin accumulation at DSB initiates the ATM-dependent one-sided loop extrusion process at either side of the break that establishes the phosphorylation of H2AX, which will spread until loop extrusion reaches a boundary element (i.e. a TAD border).

We then set to address if the increased presence of cohesin at DSBs would impact the genomic overall distribution of this protein complex following break induction. Previous work indicated a genome-wide increase of cohesin, and reinforcement of TADs following exposure to irradiation^32,33^. In agreement, we found that SCC1 enrichment was increased at cohesin-binding sites post-break induction, coinciding with an increased loop strength (Fig. 4a, top panel, compare black and grey arrows). This accumulation of SCC1 at cohesin binding sites was accompanied by a slight but significant decrease of SCC1 between cohesin peaks (“interpeaks” Fig. 4b), which would be in agreement with altered cohesin properties, such as residence time and loop extrusion rate, following DSB. Of interest, DSB-induced SCC1 accrual at loop anchors was seen within undamaged TADs but was more pronounced within damaged TADs (Fig. 4c).

**Figure 4.**
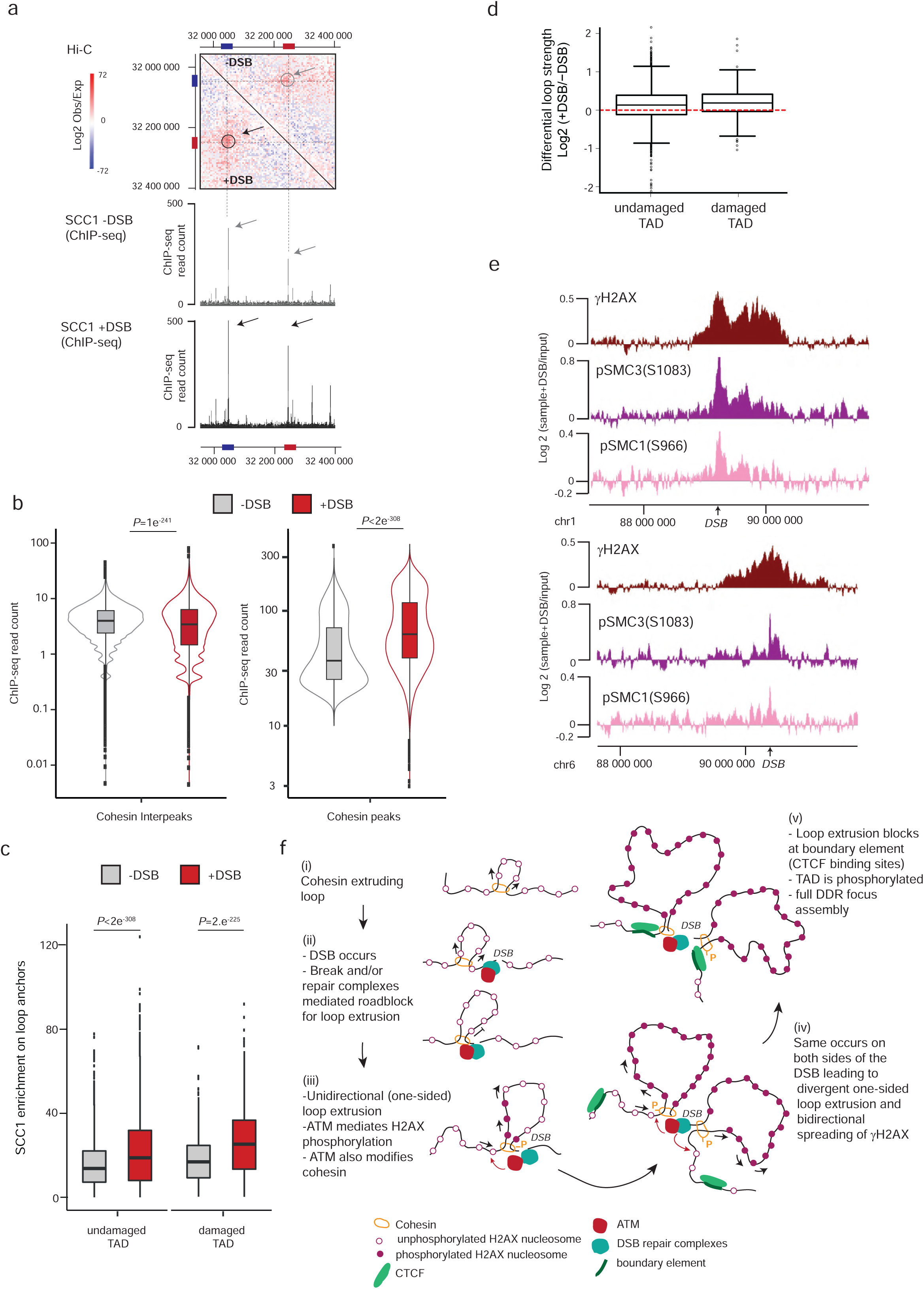
DSBs trigger modifications of cohesin biology at a genome-wide scale and accentuated in damaged TADs. **(a)** Upper panel: Contact matrix (5kb resolution) showing the Log2 (observed/expected) before or after DSB induction as indicated, on a region showing a loop on chromosome 20 and devoid of AsiSI site (no DSB). Loops anchors are circled and indicated by red and blue bars. Lower panel: Genome browser screenshot showing the SCC1 calibrated ChIP-seq on the same region before and after DSB induction as indicated. Cohesin enrichment at the loop anchors (blue and red bars) is increased after DSB (black arrows) compared to before DSB (grey arrows) in agreement with an increased loop strength (grey and black circles). **(b)** Violin plots showing the SCC1 enrichment between cohesin peaks (“inter-peaks”) (left panel) and at cohesin peaks (right panel) before and after DSB induction as indicated. P values are indicated (paired one-sided Wilcoxon test). **(c)** Boxplot showing the quantification of SCC1 recruitment on loop anchors before (grey) and after (red) DSB induction, within damaged or undamaged TADs as indicated. P values between before and after DSB are indicated (paired two-sided Wilcoxon test). The increased SCC1 enrichment on loop anchors following DSB is significantly higher in damaged TADs as compared to undamaged TADs Undamaged vs Damaged TADs, P= 3×10-38 (one-sided Wilcoxon test). **(d)** Boxplot showing the differential loops strength in undamaged or damaged TADs as indicated (see methods), computed from Hi-C data obtained before and after DSB. Undamaged TADs P= 1×10-63 (one-sided Wilcoxon test, µ=0); Damaged TADs P= 9×10-13 (one-sided Wilcoxon test, µ=0); Undamaged vs Damaged TADs, P= 0.03 (two-sided Wilcoxon test). **(e)** Genome browser screenshot showing γH2AX, phosphorylated SMC3 (p-SMC3 S1083) and phosphorylated SMC1 (p-SMC1 S966) ChIP-chip signal expressed as the log2 ratio sample/input after DSB induction and smoothed using a 500 probes span. Two damaged genomic locations are shown: on chromosome 1 (upper panel) and on chromosome 6 (lower panel). **(f)** Model: Cohesin-mediated loop extrusion ensures γH2AX establishment on the entire damaged TAD. (i) Loop extrusion constantly occurs on the genome. (ii) The occurrence of a DSB creates a roadblock for cohesin-mediated loop extrusion leading to accumulation of cohesin at the site of damage. (iii) Cohesin anchored (or blocked) at DSB continues to mediate loop extrusion in a unidirectional manner (i.e. one-sided loop extrusion, arrows). ATM, which is recruited at the immediate vicinity of the break, phosphorylates H2AX-containing nucleosomes as they are extruded. Meanwhile, cohesin are also phosphorylated by ATM, which may modify loop extrusion properties (velocity, processivity etc…) and/or cohesin turnover (chromatin loading/unloading). (iv) The same process takes place on both sides of the DSB, leading to divergent one-sided loop extrusion at either side of the break ensuring a bidirectional spreading of γH2AX. (v) Ongoing loop extrusion triggers enlargement of γH2AX modified chromatin. Loop extrusion on both sides of the DSB halt at boundary elements such as CTCF-bound loci, that demarcate TAD borders. Given that the rate of loop extrusion measured in vitro is as high as 0.5 kb per second, this process ensures that the entire damaged TAD is phosphorylated in 10-30 minutes, giving rise to a DDR focus. Whether MDC1, 53BP1, ubiquitin and H1 depletion occurs as loop extrudes or once γH2AX domain is fully established remains to be determined. NB: here the cohesin is shown as a ring encircling DNA for graphical reasons, but it is not known yet, whether and how cohesin ring entraps DNA during the loop extrusion process.

In agreement, Aggregate Peak Analysis (APA) showed that the strength of chromatin loops was significantly increased within TADs that experienced a DSB compared to undamaged TADs (Fig. 4d, Fig. S3). Thus, our data indicate a generalized increase in SCC1 occupancy and loop strength throughout the genome after DSB production in a manner that is even more reinforced within the TADs subjected to DSB. The SMC1 and SMC3 cohesin subunits have been previously reported to be phosphorylated by ATM following DSBs induction^34^ and these modifications are essential for cohesin reinforcement on the genome post irradiation^32^. Interestingly, we found that phosphorylated SMC1 (pSMC1 S966) and SMC3 (pSMC3 S1083) accumulate on entire TADs around DSBs (Fig. 4e). Altogether these data suggest that these DSB-induced, ATM-mediated cohesin modifications accumulating around DSBs, may regulate cohesin properties, such as loop extrusion velocity or chromatin unloading and translate into increased cohesin residence time at boundaries elements.

In summary, our work unambiguously shows that TADs are the template for the spreading of many DSB repair signaling events such as the phosphorylation of H2AX, the eviction of histone H1 and the accrual of 53BP1, MDC1 and ubiquitin, allowing a DSB signaling at the megabase scale. Our data are in agreement with a DSB-anchored cohesin-mediated loop extrusion model that would mediate H2AX phosphorylation (Fig. 4f). In this model, cohesin rapidly accumulates on both sides of a DSB due to a boundary function of the break itself that would naturally impair loop extrusion, or due to *de novo* loading of the cohesin at sites of damage. Divergent one-sided loop extrusion hence takes place at the DSB, which in turn allows the locally-recruited ATM to phosphorylate H2AX containing nucleosomes as the chromatin fiber is pulled by the cohesin ring. Given that current estimates of cohesin-mediated loop extrusion suggest a rate of 0.5-2 kb per second *in vitro* ^24,25^, such a mechanism would allow a rapid way to assemble DDR foci, with the entire megabase-sized chromatin domain being modified in about 10-30 min, which fits with the observed rate of γH2AX foci assembly^9^. This model would be in agreement with the recent finding that in yeast, the ATM ortholog Tel1 mediates H2A phosphorylation in a manner that agrees with a 1D sliding model rather than a 3D diffusion model ^35^. Moreover, our data also indicate that the cohesin complex itself is modified by ATM which may alter the properties of cohesin such as loop extrusion velocity, its ability to go beyond boundary element or its capability to load onto/unload from chromatin.

Recent works support a key function of genome architecture, TADs borders and loop extrusion in genome stability, including immunoglobulin loci rearrangements^36,37^ and in DSB occurrence through topoisomerase reactions^38,39^. Our study shows that genome architecture is also instrumental for the correct establishment of γH2AX and DDR foci formation, expanding the function of genome organization within TADs to the response to DNA damage. We propose that ongoing loop extrusion provides an efficient and rapid way to signal a DSB and assemble a DDR focus, while boundary elements help constraining DDR signaling to DSB-surrounding, self-interacting chromatin domains. This creates a specific repair prone chromatin compartment, that displays modified dynamics properties, which may, for example reduce search time for DNA end rejoining and homology search, and/or concentrate repair factors.

## Acknowledgments

We thank the genomics core facility of EMBL for high throughput sequencing as well as the the facilities and expertise of the high throughput sequencing core facility of the I2BC (Centre de Recherche de Gif).

Funding in G.L.’s laboratory was provided by grants from the European Research Council (ERC-2014-CoG 647344), the Agence Nationale pour la Recherche (ANR-14-CE10-0002-01 and ANR-13-BSV8-0013), the Institut National Contre le Cancer (INCA), the Ligue Nationale Contre le Cancer (LNCC). C.A is a recipient of a FRM fellowship (FRM FDT201904007941).

## Author contributions

C.A., T.C. and P.C. performed and analyzed experiments. V.R. and R.M. performed bioinformatic analyses of 4C-seq, Hi-C and ChIP-seq data sets. E.P.R. and P.E.M. provided nanoblades for CRISPR/cas9 experiments. D.N. helped to realize and analyze 4C-seq experiments. G.L. conceived experiments and G.L. and T.C. supervised experiments. G.L. wrote the manuscript. All authors commented and edited the manuscript.

## Methods

### Cell culture and treatments

DIvA (AsiSI-ER-U20S) cells were grown in Dubelcco’s modified Eagle’s medium (DMEM) supplemented with 10% SVF (InVitrogen), antibiotics and 1 µg/mL puromycin (DIvA cells) at 37 °C under a humidified atmosphere with 5% CO2. For DSB induction, cells were treated with 300nM 4OHT (Sigma, H7904) for 4 h. For ATM inhibition, cells were pretreated for 1 h with 20μM KU-55933 (Sigma, SML1109) and during subsequent 4OHT treatment. siRNA transfections were performed with a siRNA Ctrl: CAUGUCAUGUGUCACAUCU and a siRNA targeting SCC1: GGUGAAAAUGGCAUUACGG, using the cell line Nucleofector kit V (Program X-001, Amaxa) according to the manufacturer’s instructions and subsequent treatment(s) were performed 48 h later. For CRISPR/Cas9-mediated DSB induction, sgRNA (AsiSI site position: “CGCCGCGATCGCGGAATGGA” or Position further within the TAD: “GGGCCAGTCGCGGCACTCGC”) were delivered in U2OS cells using the “nanoblades” technology which relies on direct cell transduction with a viral-derived particle containing the Cas9/sgRNA ribonucleoprotein ^40,41^. Cells were analyzed 24 h after transduction.

### Immunofluorescence

DIvA cells were plated on glass coverslips and fixed with 4% paraformaldehyde during 15 min at room temperature, permeabilized with 0,5% Triton X-100 in PBS for 10 min then blocked with 3% BSA in PBS for 30min. Cells were then incubated with the primary antibody (referenced in Table S2) diluted in PBS-BSA overnight at 4°C, washed with 1X PBS and incubated with the appropriate anti-mouse or anti-rabbit secondary antibodies (conjugated to Alexa 594 or Alexa 488, Invitrogen), diluted 1:1000 in PBS-BSA, for 1h at room temperature, followed by DAPI staining. Coverslips were mounted in Citifluor (Citifluor, AF-1). Image acquisition was performed with MetaMorph on a wide-field microscope (Leica, DM6000) equipped with a camera (DR-328G-C01-SIL-505, ANDOR Technology) using 40x or 100x objectives. For the quantification, cells were acquired with a 40x objective and analyzed using Columbus software (Perkin Elmer). γH2AX foci were detected using the method D.

### Western Blot

Cells were incubated in RIPA buffer (50 mM Tris at pH 8, 150 mM NaCl, 0.5% deoxycholate, 1% NP-40, 0.1% SDS) for 20 min on ice and centrifuged at 13,000 rpm for 10 min to remove insoluble material and SDS loading buffer and reducting agent were then added to the supernatant. Protein extracts were resolved on 3%–8% NuPAGE Tris-acetate gels (Invitrogen) and transferred onto PVDF membranes (Invitrogen) according to the manufacturer’s instructions. Membranes were blocked in TBS containing 0.1% Tween 20 (Sigma, P1379) and 3% nonfat dry milk for 1h followed by an overnight incubation at 4°C with primary antibodies (referenced in Table S2). The appropriate horseradish peroxidase-coupled secondary antibodies were used to reveal the proteins (antimouse at 1:10,000 (Sigma, A2554) and antirabbit at 1: 10,000 (Sigma, A0545)) using a luminol-based enhanced chemiluminescence HRP substrate (Super Signal West Dura Extended Duration Substrate, Thermo Scientific). Pictures of the membranes were acquired with the ChemiDoc™ Touch Imaging System.

### Hi-C

Hi-C experiments were performed in DIvA cells using the Arima Hi-C kit (Arima Genomics) according to the manufacturer’s instructions. 1×10^6^ cells were used by condition and experiments were performed in duplicates. Briefly, cells were cross-linked with 2% formaldehyde for 10 min at RT, lysed and chromatin was digested with two different restriction enzymes included in the kit. Ends were filled-in in the presence of biotinylated nucleotides, followed by subsequent ligation. Ligated DNA was sonicated using the Covaris S220 to an average fragment size of 350 bp with the following parameters (Peak incident power: 140; Duty factor: 10%; Cycles per burst: 200; Treatment time: 70s). DNA was then subjected to a double-size selection to retain DNA fragments between 200 and 600 bp using Ampure XP beads (Beckman Coulter). Biotin-ligated DNA was precipitated with streptavidin-coupled magnetic beads (included in the kit). Hi-C library was prepared on beads using the NEBNext® Ultra™ II DNA Library Prep Kit for Illumina and NEBNext® Multiplex Oligos for Illumina (New England Biolabs) following instructions from the Arima Hi-C kit. The final libraries were subjected to 75 bp paired-end sequencing on a Nextseq500 platform at the EMBL Genomics core facility (Heidelberg, Germany). Hi-C reads were mapped to hg19 and processed with Juicer using default settings (https://github.com/aidenlab/juicer). Matrix-balanced Hi-C count matrices were generated at multiple resolutions: 100 kb, 50 kb, 25 kb, 10 kb and 5 kb.

### 4C-seq

The 4C-seq experiments were realized as in ^42^ with minor modifications. Briefly, 15×10^6^ DIvA cells were cross-linked with 2% formaldehyde for 10 min at RT, lysed and digested with MboI (New England Biolabs). Two or three rounds of 4 h of digestion with MboI were necessary. Digested DNA was then ligated with a T4 DNA ligase (HC) (Promega), purified and digested with NlaIII overnight (New England Biolabs). After a second ligation step, DNA was purified before proceeding to the library preparation. For DNA purification steps, AMPure XP beads (Beckman Coulter) were used diluted at 1:10 in 20% PEG solution (PEG 8000 (Sigma) 20%, 2.5M NaCl, Tween 20 20%, Tris pH8 10mM, EDTA 1mM). For 4C-seq library preparation, 800ng-900ng of 4C-seq template was amplified using 16 individual PCR reactions with inverse primers (PAGE-purified) including the Illumina adapter sequences and a unique index for each condition (primers in Table S1). Libraries were purified with the QIAquick PCR Purification Kit (Qiagen), pooled and subjected to 75 bp single-end sequencing on a Nextseq500 platform at the I2BC Next Generation Sequencing Core Facility (Gif-sur-Yvette, France). Each sample was then demultiplexed using a specific python script from the FourCSeq R package^43^ thus assigning each read to a specific viewpoint based to its primer sequence into separate fastQ files. bwa mem was then used for mapping and samtools for sorting and indexing. A custom R script (https://github.com/bbcf/bbcfutils/blob/master/R/smoothData.R) ^44^ was used to build the coverage file in bedGraph format, to normalize using the average coverage and to exclude the nearest region from each viewpoint (viewpoint-containing restriction fragment and the two adjacent restriction fragments). Then the bedGraph file was converted in a BigWig file using bedGraphToBigWig program from UCSC.

### ChIP-seq and ChIP-chip

For Fig. 1 a, Ubiquitin, H1, γH2AX and 53BP1 ChIP-seq data were retrieved from ^5^. ChIP experiments of P-ATM, MDC1 and γH2AX were performed in DIvA cells as in ^10^ with 200µg of chromatin per immunoprecipitation. The antibodies used are detailed in Table S2. Prior to library preparation, samples from multiple ChIP experiments were pooled and sonicated for 15 cycles (30 sec ON, 30 sec OFF, high setting) with a Bioruptor (Diagenode) then concentrated with a vacuum concentrator (Eppendorf). Sequencing libraries were prepared by using 10 ng of purified DNA (averaged size 250-300 bp) with the NEBNext® Ultra™ II Library Prep Kit for Illumina (New England Biolabs) using the application note for “Low input ChIP-seq.”, and subjected to 75 bp single-end sequencing on a Nextseq500 platform at the EMBL Genomics core facility (Heidelberg, Germany).

For the calibrated ChIP-seq of SCC1, we used a spike-in method ^45^. Briefly, cross-linked cells were first lysed 10min at 4°C in 500 µL of the Lysis Buffer 1 (10mM Tris pH8, 10mM NaCl, 0.5% NP-40, Complete protease inhibitor (Sigma)) then 10min at 4°C in the Lysis buffer 2 (50mM Tris pH8, 10mM EDTA, 0.5% NP-40, Complete protease inhibitor (Sigma)) and subsequently sonicated in 15ml conical tubes with a Bioruptor Pico (Diagenode) in the presence of 800 mg of sonication beads (20 cycles of 30 sec ON/30 sec OFF) to an average fragment size of 250pb. Prior to immunoprecipitation with SCC1 antibody, 20% of chromatin from mouse ES cells (40µg) was added to chromatin prepared from treated or untreated human DIvA cells (200µg). Sequencing libraries were prepared from IP and input samples using the NEBNext® Ultra™ II Library Prep Kit for Illumina and subjected to 75 bp single-end sequencing on a Nextseq500 platform at the EMBL Genomics core facility (Heidelberg, Germany). First, SCC1 was aligned on the mouse genome (mm10) with bwa in order to only map the reads used as a reference for the normalization (spike-in). Remaining unmapped reads were re-converted into a fastQ file using bam2fastq and mapped to the human genome (hg19) using bwa. Samtools was used for sorting and indexing, and reads mapped to the mouse genome were used as a normalization factor, as in ^45^ and using the following formula.

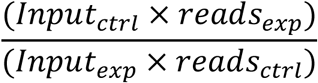

where *Input*_*ctrl*_ is the total number of reads mapped in ES Input (Mouse) and *Input*_*exp*_ the total number of reads in DIvA Input. *reads*_*ctrl*_ and *reads*_*exp*_ were respectively the number of reads from immunoprecipitated samples mapped on the mm10 genome and hg19 genome.

For the ChIP-chip experiments, the immunoprecipited samples of γH2AX, SMC1 (pSMC1 S966) or SMC3 (p-SMC3 S1083) and input samples were amplified as in ^10^, labeled, and hybridized on Affymetrix tiling arrays covering human chromosomes 1 and 6 (at the Genotoul GeT-biopuces facility (Toulouse, France), or the EMBL genomic core platform). Scanned array data were normalized using Tiling Affymetrix Software (TAS) (quantile normalization, scale set to 500) and analyzed as described in ^10,12^ and converted to .wig files using R/Bioconductor software, when necessary, for visualization using the Integrated Genome Browser (bioviz.org).

### Hi-C, 4C-seq and ChIP-seq analyses

#### Hi-C heatmaps

Hi-C heatmaps screenshots were generated using Juicebox stand-alone program (https://github.com/aidenlab/Juicebox/wiki/Download). To build the average heatmaps, sub-matrix for *cis* interactions around DSBs were extracted using Juicer. Log2(ratio) after vs before DSB were computed, using both Hi-C replicates, and averaged for each bin of the final matrix.

#### Insulation score and TAD calling

Insulation score was computed using Hi-C matrices at 50 kb resolution with matrix2insulation.pl (https://github.com/dekkerlab/crane-nature-2015). As parameters, we used is=800000 and ids=100000. TADs were called using Hi-C matrices at 50 kb resolution with TopDom R package and window size parameter of 10 (https://github.com/HenrikBengtsson/TopDom). In order to filter out very weak TAD borders (corresponding to sub-TAD borders), we filtered TAD borders with an insulation score below a threshold of -0.05.

#### Loops anchors and APA

Loops were called using Juicer Tools HiCCUPS program at 10 kb and 25 kb resolutions (https://github.com/aidenlab/juicer/wiki/HiCCUPS). Aggregate Peak Analysis (APA) was done using Juicer Tools APA program at 10 kb resolution (https://github.com/aidenlab/juicer/wiki/APA). For Fig 2d, 525 loops were retrieved between the 174 best cleaved DSBs and nearby loop anchors (<1Mb) Fold change between Signal (central pixel) and background (upper left corner 5×5 pixels) was computed. For Fig. S3, 1226 and 6577 loops in damaged (174 damaged TADs) and undamaged TADs were retrieved respectively. Fold change between Signal (central pixel) and background (lower left corner 5×5 pixels) was computed. APA heatmaps were reprocessed using ggplot2 in order to display counts at the same color scale between -DSB and +DSB conditions. For Fig. 4d, APA were generated for loops filtered on their size (<200kb) and around the best 80 cleaved DSBs. Loop strength was extracted from APA files enhancement.txt corresponding to enrichment fold-change (Peak to Mean, P2M). Differential loop strength was the log-ratio of two condition loop strengths (+DSB/-DSB).

#### SCC1 ChIP-seq analyses

For Fig. 4b, peaks were identified using MACS2 program with callpeak algorithm, with default setting, using Input as control and the SCC1 ChIP-seq data before break induction as sample. Before breaks, 46184 peaks were identified, with a median and a mean size of 628 and 742 respectively. An equivalent number of cohesin unbound loci were picked in the inter-peak fragment, with an equivalent size distribution.

#### 4C-seq

For differential analyses of the 4C-seq data, the log2 ratio between two bam files was computed using bamCompare from deeptools, with the parameters --binSize=50 and --operation=log2. Each boxplot represents the mean of the 4C-seq ratio on 1 MB around each viewpoint.

## Data Availability

High throughput sequencing data have been deposited to Array Express under accession number E-MTAB-XXX. Other data and source codes are available upon request.

---

**Supplementary Figure 1.**
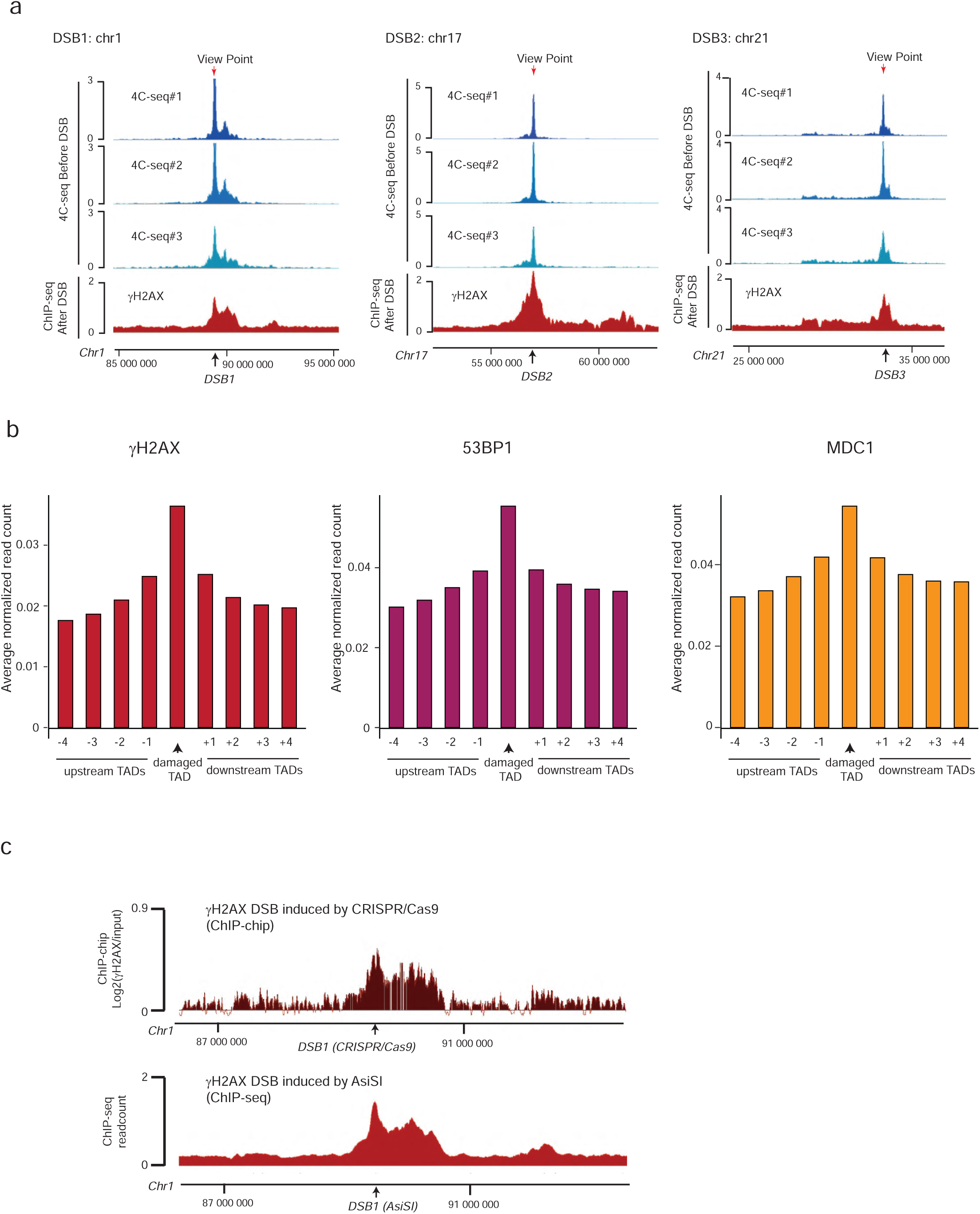
Related to figure 1. **(a)** Genome browser screenshots representing the 4C-seq signal before DSB induction (-DSB) obtained for three independent biological replicates and the γH2AX ChIP-seq signal after DSB induction for different viewpoints localized at three AsiSI sites or a Control region. ChIP-seq data are smoothed using 100 kb span, 4C-seq using a 50 kb span. The AsiSI sites are indicated by arrows. **(b)** ChIP-seq signal for γH2AX (left), 53BP1 (middle) and MDC1 (right) was computed within the damaged TAD and neighboring TADs for the 174 best cleaved DSBs in DIvA cells.**(c)** Genome browser screenshot of the chromosome 1 showing the γH2AX distribution around a DSB induced by CRISPR/Cas9 (upper panel, ChIP-chip, expressed as the log2 sample/input) and by AsiSI at the same position (lower panel, ChIP-seq).

**Supplementary Figure 2.**
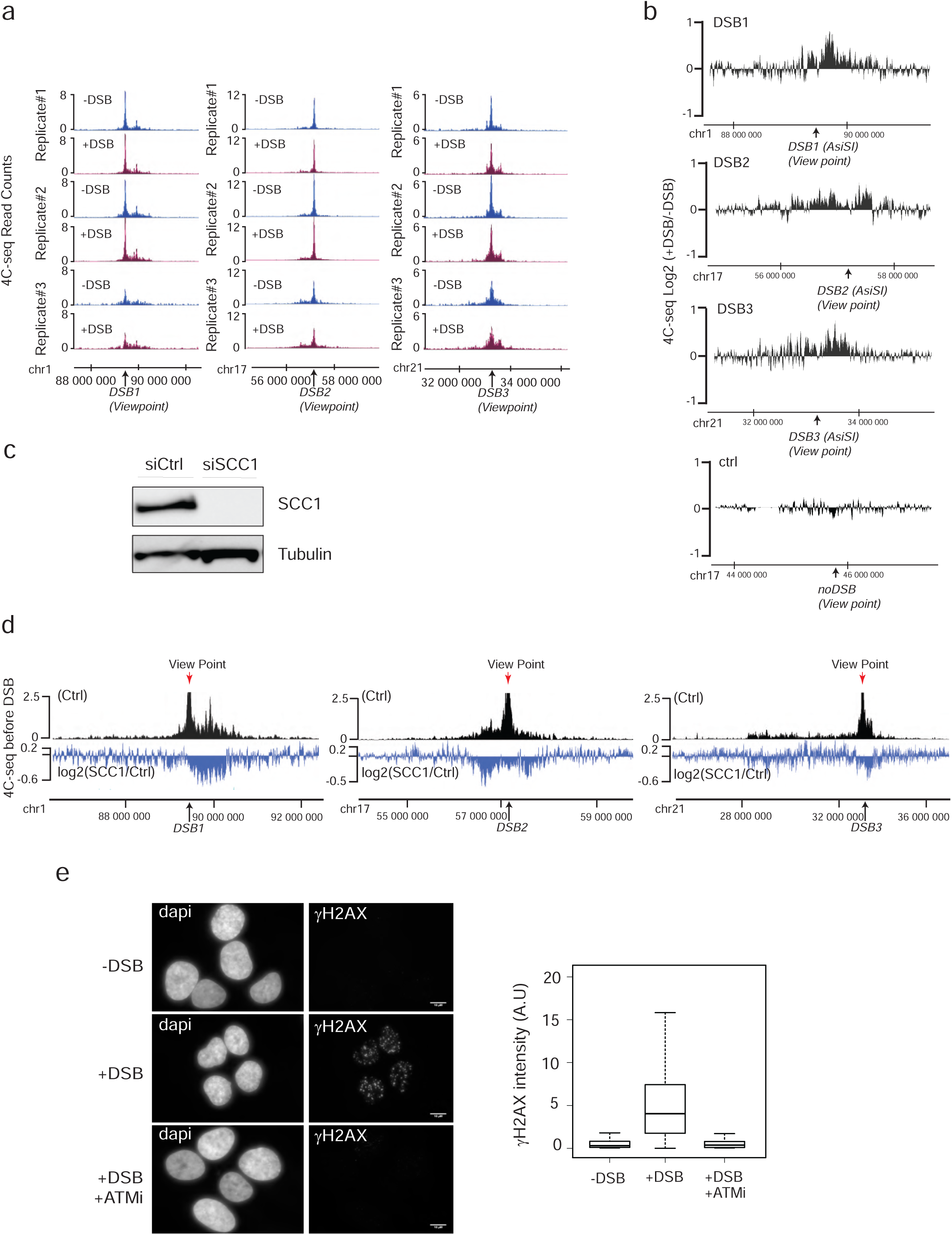
Related to figure 3. **(a)** Genome browser screenshots representing the 4C-seq signal (smoothed using a 10 kb span) before and after DSB induction as indicated, obtained for three biological replicates using viewpoints localized at three DSB sites. The DSBs are indicated by arrows. **(b)** Differential 4C-seq (Log2 +DSB/-DSB) is shown for three viewpoints located at DSB sites (viewpoints 1 to 3) and on a Ctrl region (viewpoint 4). **(c)** Western Blot showing the depletion of SCC1 by siRNA. **(d)** Genome browser screenshots showing the differential (Log2) 4C-seq signal in SCC1 siRNA treated cells versus control siRNA-treated cells (in undamaged conditions) for three viewpoints. **(e)** Left: Immunofluorescence experiment showing γH2AX and DAPI staining before and after DSB induction with or without ATM inhibitor as indicated. Right: Quantification of γH2AX intensity (expressed in A.U: Arbitrary Unit) in the above conditions. One representative experiment is shown.

**Supplementary Figure 3.**
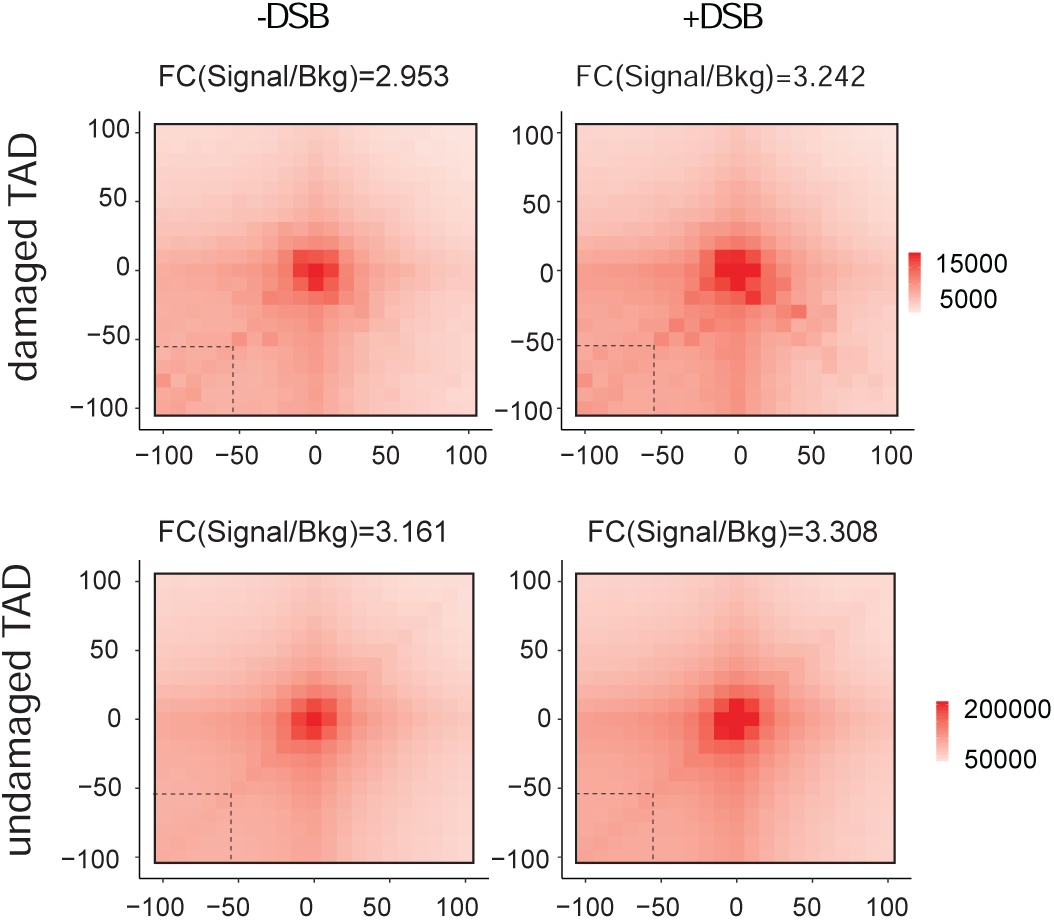
Related to figure 4. Aggregate Peak Analysis (APA) plot on a 200 kb window (10 kb resolution) before (-DSB) and after DSB induction (+DSB) calculated for all loops anchors, in damaged and undamaged TAD as indicated. The fold change between the signal (central pixel) and the background (lower left corner 5×5 pixels) is indicated.

**Supplementary Table 1:**
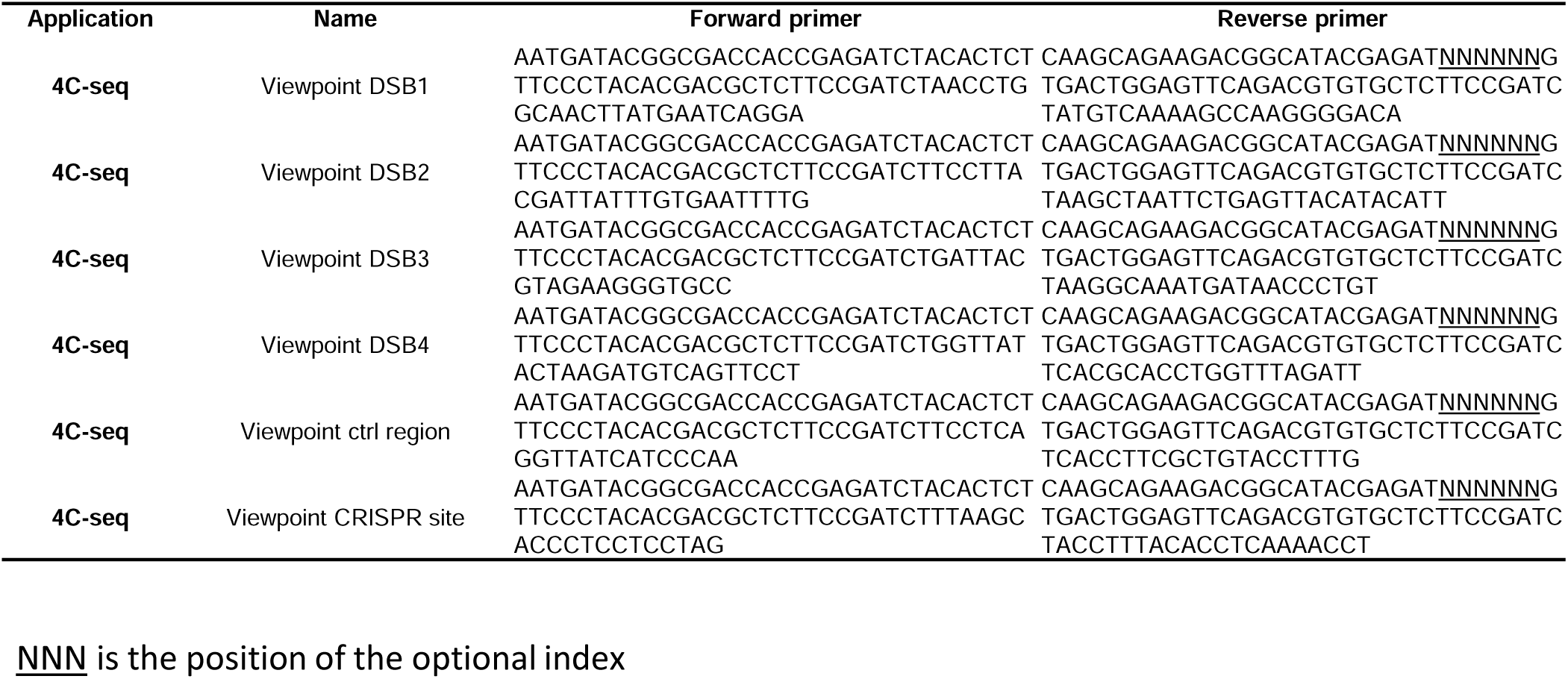
Primer sequences

**Supplementary Table 2:** Antibodies

| Target | Application | Reference | Quantity |
| --- | --- | --- | --- |
| γH2AX (S139) | ChIP | Merck Millipore 07-164 | 2 µg |
| γH2AX (S139) | IF | Merck Millipore 05-636 (clone JBW301) | 1/1000 |
| P-ATM (S1981) | ChIP | Abcam ab81292 | 2 µg |
| MDC1 | ChIP | Abcam ab11171 | 3 µg |
| SCC1 | ChIP | Abcam ab992 | 4 µg |
| SCC1 | Western Blot | Abcam ab992 | 1/500 |
| SMC1 (P966) | ChIP | Abcam ab81306 (clone EP2858Y) | 2 µL |
| SMC3 (P1083) | ChIP | Bethyl Laboratories A300-480A | 2 µg |

## References

1. Clouaire, T., Marnef, A. & Legube, G. Taming Tricky DSBs: ATM on duty. DNA Repair (Amst.) 56, 84–91 (2017).

2. McCord, R. P., Kaplan, N. & Giorgetti, L. Chromosome conformation capture and beyond: toward an integrative view of chromosome structure and function. Mol. Cell (2020). doi:10.1016/j.molcel.2019.12.021

3. Arnould, C. & Legube, G. The Secret Life of Chromosome Loops upon DNA Double-Strand Break. J. Mol. Biol. (2019). doi:10.1016/j.jmb.2019.07.036

4. Rogakou, E. P., Boon, C., Redon, C. & Bonner, W. M. Megabase chromatin domains involved in DNA double-strand breaks in vivo. J. Cell Biol. 146, 905–916 (1999).

5. Clouaire, T. et al. Comprehensive Mapping of Histone Modifications at DNA Double-Strand Breaks Deciphers Repair Pathway Chromatin Signatures. Mol. Cell 72, 250–262.e6 (2018).

6. Stewart, G. S., Wang, B., Bignell, C. R., Taylor, A. M. R. & Elledge, S. J. MDC1 is a mediator of the mammalian DNA damage checkpoint. Nature 421, 961–966 (2003).

7. Caron, P. et al. Cohesin protects genes against γH2AX Induced by DNA double-strand breaks. PLoS Genet. 8, e1002460 (2012).

8. Natale, F. et al. Identification of the elementary structural units of the DNA damage response. Nat. Commun. 8, 15760 (2017).

9. Ochs, F. et al. Stabilization of chromatin topology safeguards genome integrity. Nature 574, 571–574 (2019).

10. Iacovoni, J. S. et al. High-resolution profiling of gammaH2AX around DNA double strand breaks in the mammalian genome. EMBO J. 29, 1446–1457 (2010).

11. Chang, L.-H., Ghosh, S. & Noordermeer, D. Tads and their borders: free movement or building a wall? J. Mol. Biol. (2019). doi:10.1016/j.jmb.2019.11.025

12. Caron, P. et al. Non-redundant Functions of ATM and DNA-PKcs in Response to DNA Double-Strand Breaks. Cell Rep. 13, 1598–1609 (2015).

13. Schwarzer, W. et al. Two independent modes of chromatin organization revealed by cohesin removal. Nature 551, 51–56 (2017).

14. Rao, S. S. P. et al. Cohesin loss eliminates all loop domains. Cell 171, 305–320.e24 (2017).

15. Gelot, C. et al. The Cohesin Complex Prevents the End Joining of Distant DNA Double-Strand Ends. Mol. Cell 61, 15–26 (2016).

16. Meisenberg, C. et al. Repression of Transcription at DNA Breaks Requires Cohesin throughout Interphase and Prevents Genome Instability. Mol. Cell 73, 212–223.e7 (2019).

17. Potts, P. R., Porteus, M. H. & Yu, H. Human SMC5/6 complex promotes sister chromatid homologous recombination by recruiting the SMC1/3 cohesin complex to double-strand breaks. EMBO J. 25, 3377–3388 (2006).

18. Unal, E. et al. DNA damage response pathway uses histone modification to assemble a double-strand break-specific cohesin domain. Mol. Cell 16, 991–1002 (2004).

19. Unal, E., Heidinger-Pauli, J. M. & Koshland, D. DNA double-strand breaks trigger genome-wide sister-chromatid cohesion through Eco1 (Ctf7). Science 317, 245–248 (2007).

20. Ström, L. et al. Postreplicative formation of cohesion is required for repair and induced by a single DNA break. Science 317, 242–245 (2007).

21. Ström, L., Lindroos, H. B., Shirahige, K. & Sjögren, C. Postreplicative recruitment of cohesin to double-strand breaks is required for DNA repair. Mol. Cell 16, 1003–1015 (2004).

22. Bauerschmidt, C. et al. Cohesin promotes the repair of ionizing radiation-induced DNA double-strand breaks in replicated chromatin. Nucleic Acids Res. 38, 477–487 (2010).

23. Covo, S., Westmoreland, J. W., Gordenin, D. A. & Resnick, M. A. Cohesin Is limiting for the suppression of DNA damage-induced recombination between homologous chromosomes. PLoS Genet. 6, e1001006 (2010).

24. Davidson, I. F. et al. DNA loop extrusion by human cohesin. Science 366, 1338–1345 (2019).

25. Kim, Y., Shi, Z., Zhang, H., Finkelstein, I. J. & Yu, H. Human cohesin compacts DNA by loop extrusion. Science 366, 1345–1349 (2019).

26. Ganji, M. et al. Real-time imaging of DNA loop extrusion by condensin. Science 360, 102–105 (2018).

27. Vian, L. et al. The energetics and physiological impact of cohesin extrusion. Cell 173, 1165–1178.e20 (2018).

28. Schmitt, A. D. et al. A compendium of chromatin contact maps reveals spatially active regions in the human genome. Cell Rep. 17, 2042–2059 (2016).

29. Mirny, L. A., Imakaev, M. & Abdennur, N. Two major mechanisms of chromosome organization. Curr. Opin. Cell Biol. 58, 142–152 (2019).

30. Aymard, F. et al. Transcriptionally active chromatin recruits homologous recombination at DNA double-strand breaks. Nat. Struct. Mol. Biol. 21, 366–374 (2014).

31. Bot, C. et al. Independent mechanisms recruit the cohesin loader protein NIPBL to sites of DNA damage. J. Cell Sci. 130, 1134–1146 (2017).

32. Kim, B.-J. et al. Genome-wide reinforcement of cohesin binding at pre-existing cohesin sites in response to ionizing radiation in human cells. J. Biol. Chem. 285, 22784–22792 (2010).

33. Sanders, J. T. et al. Radiation-Induced DNA Damage and Repair Effects on 3D Genome Organization. BioRxiv (2019). doi:10.1101/740704

34. Kim, S.-T., Xu, B. & Kastan, M. B. Involvement of the cohesin protein, Smc1, in Atm-dependent and independent responses to DNA damage. Genes Dev. 16, 560–570 (2002).

35. Li, K., Bronk, G., Kondev, J. & Haber, J. E. Yeast ATM and ATR use different mechanisms to spread histone H2A phosphorylation around a DNA double-strand break. BioRxiv (2019). doi:10.1101/2019.12.17.877266

36. Zhang, Y. et al. The fundamental role of chromatin loop extrusion in physiological V(D)J recombination. Nature 573, 600–604 (2019).

37. Zhang, X. et al. Fundamental roles of chromatin loop extrusion in antibody class switching. Nature 575, 385–389 (2019).

38. Gothe, H. J. et al. Spatial chromosome folding and active transcription drive DNA fragility and formation of oncogenic MLL translocations. Mol. Cell 75, 267–283.e12 (2019).

39. Canela, A. et al. Topoisomerase II-Induced Chromosome Breakage and Translocation Is Determined by Chromosome Architecture and Transcriptional Activity. Mol. Cell 75, 252–266.e8 (2019).

40. Mangeot, P. E. et al. Genome editing in primary cells and in vivo using viral-derived Nanoblades loaded with Cas9-sgRNA ribonucleoproteins. Nat. Commun. 10, 45 (2019).

41. Marnef, A. et al. A cohesin/HUSH- and LINC-dependent pathway controls ribosomal DNA double-strand break repair. Genes Dev. 33, 1175–1190 (2019).

42. Matelot, M. & Noordermeer, D. Determination of High-Resolution 3D Chromatin Organization Using Circular Chromosome Conformation Capture (4C-seq). Methods Mol. Biol. 1480, 223–241 (2016).

43. Klein, F. A. et al. FourCSeq: analysis of 4C sequencing data. Bioinformatics 31, 3085–3091 (2015).

44. David, F. P. A. et al. HTSstation: a web application and open-access libraries for high-throughput sequencing data analysis. PLoS One 9, e85879 (2014).

45. Kojic, A. et al. Distinct roles of cohesin-SA1 and cohesin-SA2 in 3D chromosome organization. Nat. Struct. Mol. Biol. 25, 496–504 (2018).

